# KwARG: Parsimonious Reconstruction of Ancestral Recombination Graphs with Recurrent Mutation

**DOI:** 10.1101/2020.12.17.423233

**Authors:** Anastasia Ignatieva, Rune B. Lyngsø, Paul A. Jenkins, Jotun Hein

## Abstract

The reconstruction of possible histories given a sample of genetic data in the presence of recombination and recurrent mutation is a challenging problem, but can provide key insights into the evolution of a population. We present KwARG, which implements a parsimony-based greedy heuristic algorithm for finding plausible genealogical histories (ancestral recombination graphs) that are minimal or near-minimal in the number of posited recombination and mutation events. Given an input dataset of aligned sequences, KwARG outputs a list of possible candidate solutions, each comprising a list of mutation and recombination events that could have generated the dataset; the relative proportion of recombinations and recurrent mutations in a solution can be controlled via specifying a set of ‘cost’ parameters. We demonstrate that the algorithm performs well when compared against existing methods. The software is made available on GitHub.

## 1. Introduction

For many species, the evolution of genetic variation within a population is driven by the processes of mutation and recombination in addition to genetic drift. A typical mutation affects the genome at a single position, and may or may not spread through subsequent generations by inheritance. Recombination, on the other hand, occurs when a new haplotype is created as a mixture of genetic material from two different sources, which can drive evolution at a much faster rate. The detection of recombination is an important problem which can provide crucial scientific insights, for instance in understanding the potential for rapid changes in pathogenic properties within viral populations (Simon-Loriere and Holmes, 2011).

Consider a population evolving through the replication, mutation, and recombination of genetic material within individuals, emerging from a common origin and living through multiple generations until the present day. In general, the history of shared ancestry, mutation, and recombination events are not observed, and must be inferred from a sample of genetic data obtained from the present-day population. Crossover recombination can occur anywhere along a sequence, and the breakpoint position is also unobserved. This article focuses on methods for reconstructing possible histories of such a sample, in the form of *ancestral recombination graphs (ARGs)* — networks of evolution connecting the sampled individuals to shared ancestors in the past through coalescence, mutation, and crossover recombination events; an example is illustrated in Figure 1. This is a very important but challenging problem, as many possible histories might have generated a given sample. Moreover, recombination can be undetectable unless mutations appear on specific branches of the genealogy (Hein et al., 2004, Section 5.11), and recombination events can produce patterns in the data that are indistinguishable from the effects of *recurrent mutation* (McVean et al., 2002); that is, two or more mutation events in a genealogical history that affect the same locus.

**Figure 1.**
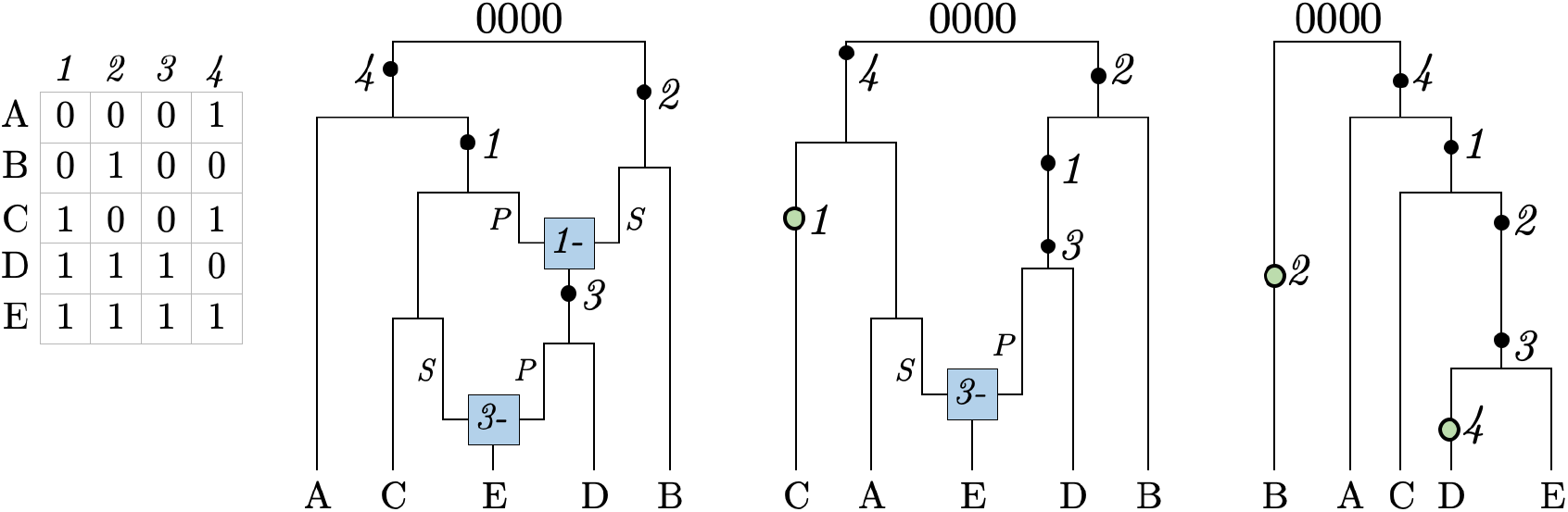
Three examples of ARGs. The dataset is shown on the left in binary format, with 0’s and 1’s corresponding to the ancestral and mutant state at each site, respectively. Mutation events are shown as black dots and labelled by the site they affect; green filled circle corresponds to a recurrent mutation. Recombination nodes (in blue) are labelled with the recombination breakpoint; material to the right (left) of the breakpoint is inherited from the parent connected by the edge labelled *S* (*P*) for “suffix” (“prefix”).

Parsimony is a widely-used approach focused on finding possible histories which minimise the number of recombinations and recurrent mutations. This does not necessarily describe the most biologically plausible version of events, but produces a useful lower bound on the complexity of the evolutionary pathway that might have generated the given dataset. Beyond specifying the types of events that are allowed, parsimony does not require assuming a particular generative model; the approach focuses on sequences of events that can generate the observed dataset, disregarding the timing and prior rate of these events.

Previous work on reconstructing histories using parsimony has tackled recombination and recurrent mutation separately. Algorithms for reconstructing minimal ARGs generally make the *infinite sites assumption*, which allows at most one mutation to have occurred at each site of the genome, thus precluding recurrent mutation events, and the goal is to calculate the minimum number of crossover recombinations required to explain a dataset, denoted *R*_*min*_. Even with this constraint, the problem is NP-hard (Wang et al., 2001); exact algorithms are practical only for small datasets (Hein, 1990; Lyngsø et al., 2005), and general methods rely on heuristic approximations (Hein, 1993; Song et al., 2005; Minichiello and Durbin, 2006; Parida et al., 2008; Thao and Vinh, 2019). Alternatively, one can assume the absence of recombination and seek to calculate the minimum number of recurrent mutations required, denoted *P*_*min*_. In this case, reconstruction of maximum parsimony trees is also NP-hard (Foulds and Graham, 1982); likewise, methods can only handle small datasets or are based on heuristics (Semple and Steel, 2003, Section 5.4).

Parsimony contrasts with the alternative approach of model-based inference, which requires the specification of a generative model and focuses on the estimation of mutation and recombination rates as model parameters. Model-based inference requires integrating over the space of possible histories, which is generally intractable; methods rely on MCMC (e.g. Rasmussen et al., 2014) or importance sampling (e.g. Jenkins and Griffiths, 2011), but the problem remains computationally difficult. If the presence of recombination is certain and reasonable models of population dynamics are available, model-based approaches may be more suitable and result in more powerful inference. However, specifying a realistic model can be prohibitively difficult, for instance when modelling viral evolution over a transmission network, where factors such as geographical structure, social clustering, and the impact of interventions are important but difficult to incorporate. In this case, model-based inference can provide misleading results, with poor quantification of uncertainty due to model misspecification, while parsimony-based methods can enable testing for the presence of recombination where this is not certain (Bruen et al., 2006). We note also the existence of numerous other methods for inference of recombination (Robertson et al., 2006) which do not explicitly reconstruct ARGs.

KwARG (“quick ARG”) is a software tool, written in C, which implements a greedy heuristic-based parsimony algorithm for reconstructing histories that are minimal or near-minimal in the number of posited recombination and mutation events. The algorithm starts with the input dataset and generates plausible histories backwards in time, adding coalescence, mutation, recombination, and recurrent mutation events to reduce the dataset until the common ancestor is reached. By tuning a set of cost parameters for each event type, KwARG can find solutions consisting only of recombinations (giving an upper bound on *R*_*min*_), only of recurrent mutations (giving an upper bound on *P*_*min*_), or a combination of both event types. KwARG handles both the ‘infinite sites’ and ‘maximum parsimony’ scenarios, as well as interpolating between these two cases by allowing recombinations as well as recurrent mutations and sequencing errors, which is not offered by existing methods. This is illustrated in Figure 1: KwARG finds all three types of solution for the given dataset. KwARG shows excellent performance when benchmarked against exact methods on small datasets, and outperforms existing heuristic-based methods on large, more complex datasets while maintaining computational efficiency. KwARG source code and executables are made freely available on GitHub at https://github.com/a-ignatieva/kwarg, along with documentation and usage examples.

The paper is structured as follows. Details of the algorithm underlying KwARG are given in Section 2, with an explanation of the required inputs and expected outputs. In Section 3, the performance of KwARG on simulated data is benchmarked against exact methods and existing programs. An application of KwARG to a widely studied *Drosophila melanogaster* dataset (Kreitman, 1983) is described in Section 4. Discussion follows in Section 5.

## 2. Technical details

Consider a sample of genetic data, where the allele at each site can be denoted 0 or 1. We do not make the infinite sites assumption, so that each site can undergo multiple mutation events. However, we do assume that mutations correspond to transitions between exactly two possible states, excluding for instance triallelic sites.

### 2.1. Input

KwARG accepts data in the form of a binary matrix, or a multiple alignment in nucleotide or amino acid format. The sequence and site labels can be provided if desired. It is possible to specify a root sequence, or leave this to be determined. The presence of missing data is permitted; regardless of the type of input, the data is converted to a binary matrix 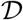, with entries ‘★’ denoting missing entries or material that is not ancestral to the sample.

### 2.2. Methods

Under the infinite sites assumption, at most one mutation is allowed to have occurred per site. If any two columns contain all four of the configurations 00, 01, 10, 11, then the data could not have been generated only through replication and mutation, and there must have been at least one recombination event between the two corresponding sites. This is the four gamete test (Hudson and Kaplan, 1985), and the two sites are said to be *incompatible*. When recurrent mutations are allowed, the incompatibility could likewise have been generated through multiple mutations affecting the same site (McVean et al., 2002).

KwARG reconstructs the history of a sample backwards in time, by starting with the data matrix 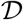 and performing row and column operations corresponding to coalescence, mutation, and recombination events, until only one ancestral sequence remains. By reversing the order of the steps, a forward-in-time history is obtained, showing how the population evolved from the ancestor to the present sample. When a choice can be made between multiple possible events, a neighbourhood of candidate ancestral states is constructed, using the same general method as that employed in the program Beagle (Lyngsø et al., 2005). A backwards-in-time approach has also been implemented in the programs SHRUB (Song et al., 2005), Margarita (Minichiello and Durbin, 2006) and GAMARG (Thao and Vinh, 2019), all of which adopt the infinite sites assumption but use different criteria for choosing amongst possible recombination events.

#### 2.2.1. Construction of a history

For convenience, assume that the all-zero sequence is specified as the root, and 0 (1) entries of 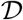 correspond to ancestral (mutated) sites. Suppose 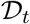 is the data matrix obtained after *t*−1 iterations of the algorithm. At the beginning of the *t*-th step, KwARG first reduces 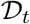, by repeatedly applying the ‘Clean’ algorithm (Song and Hein, 2003) through:

- deleting uninformative columns (consisting of all 0’s);
- deleting columns containing only one 1 (corresponding to “undoing” a mutation present in only one sequence);
- deleting a row if it agrees with another row (corresponding to a coalescence event);
- deleting a column if it agrees with an adjacent column.

Two rows (columns) *agree* if they are equal at all positions where both rows (columns) contain ancestral material, and the sites (sequences) carrying ancestral material in one are a subset of the sites (sequences) carrying ancestral material in the other.

A run of the ‘Clean’ algorithm repeatedly applies these steps to 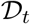, terminating when no further reduction is possible. Suppose the resulting data matrix is 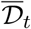. KwARG then constructs a neighbour-hood 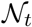 of candidate next states, each one obtained through one of the following operations:

- Pick a row and split it into two at a possible recombination point. Only a subset of possible recombining sequences and breakpoints needs to be considered; see Lyngsø et al. (2005, Section 3.3) for a detailed explanation.
- Remove a recurrent mutation, by selecting a column and changing a 0 entry to 1, or a 1 entry to 0. This is the event type that is disallowed by algorithms applying the infinite sites assumption.

Suppose a neighbourhood 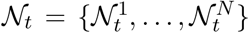 is formed, consisting of all possible states that can be reached from 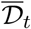 through applying one of these operations. Then the reduced neighbourhood 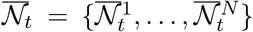 is formed by applying ‘Clean’ to each state in turn. Each state 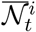 is then assigned a score 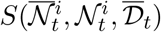 combining (i) the cost 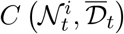, defined below, of reaching the configuration 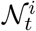 from 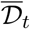, (ii) a measure 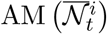 of the complexity of the resulting data matrix 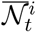, and (iii) a lower bound 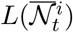 on the remaining number of recombination and recurrent mutation events still required to reach the ancestral sequence from 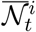. Finally, a state is selected, say 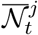, based on its score, and we set 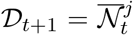. The process of reducing the dataset followed by constructing a neighbourhood and choosing the best move is repeated, until all incompatibilities are resolved and the root sequence is reached. Pseudocode for the ‘Clean’ algorithm and KwARG is given in Appendix A.

The construction of a history for the dataset given in Figure 1 is illustrated in Figure 2. The first step corresponds to the construction of a neighbourhood, two of the states 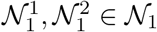 are pictured. Then, the ‘Clean’ algorithm is applied to each state in the neighbourhood (illustrated as a series of steps following blue arrows). From the resulting reduced neighbourhood 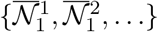, the state 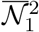 is selected; the other illustrated path is abandoned. This process is repeated until all incompatibilities are resolved and the empty state is reached. Following the path of selected moves in this figure left-to-right corresponds to the events encountered when traversing the leftmost ARG in Figure 1 from the bottom up. If instead the state 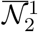 were selected at the second step of the algorithm, the resulting path would correspond to the ARG in the centre of Figure 1.

**Figure 2.**
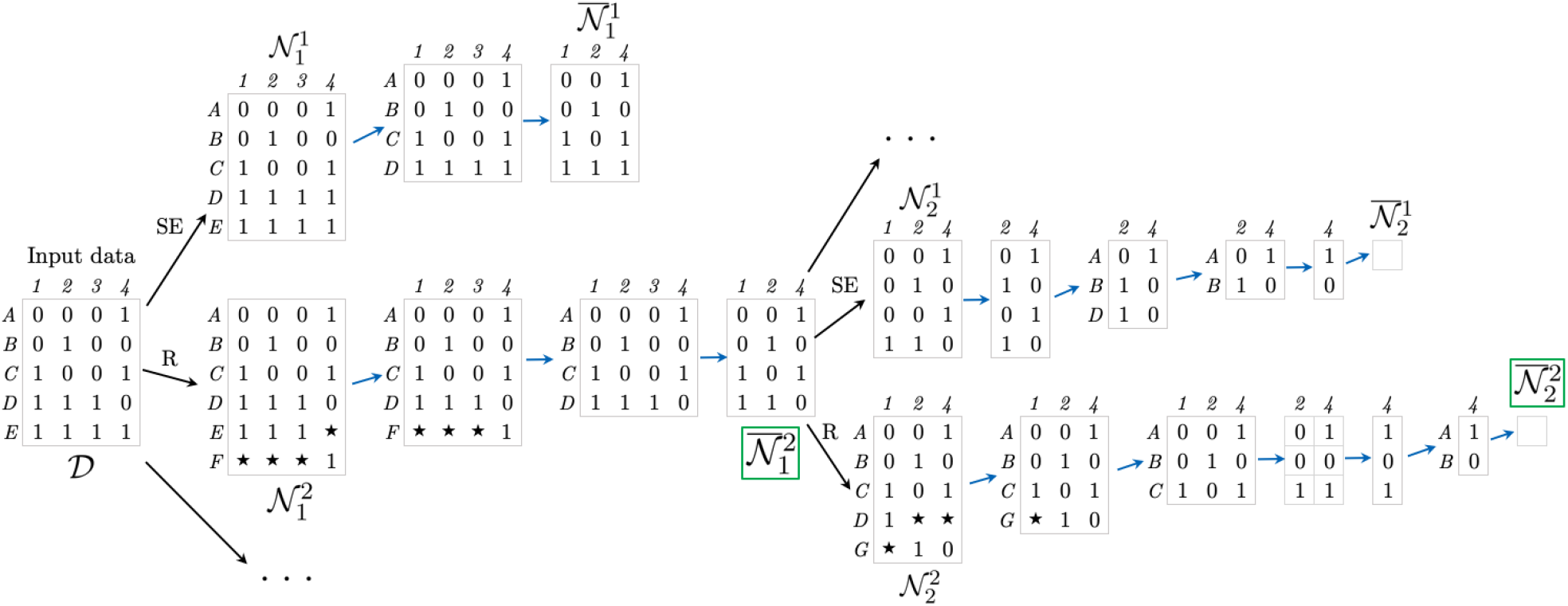
Example of a reconstructed history for the dataset in Figure 1. Stars ‘★’ denote non-ancestral material. SE: recurrent mutation occurring on a terminal branch of the ARG. R: recombination event. A sequence of blue arrows corresponds to one application of the ‘Clean’ algorithm. Green boxes highlight the selected states.

#### 2.2.2. Score

When considering which next step to take, more informed choices can be made by considering not just the cost of the step, but also the complexity of the configuration it leads to. This is the principle behind the A* algorithm (Hart et al., 1968), using a heuristic estimate of remaining distance to guide the choice of the next node to expand. KwARG applies the same principle in a greedy fashion, following a path of locally optimal choices in an attempt to find a minimal history.

The score implemented in KwARG is

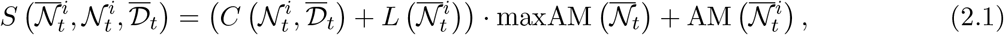

where

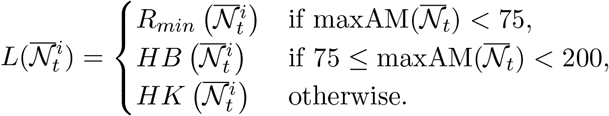

Here, 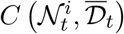 denotes the cost of the corresponding event, defined in Section 2.2.3; 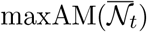 denotes the maximum amount of ancestral material seen in any of the states in 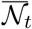, and 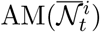 gives the amount of ancestral material in state 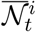. Incorporating a measure of the amount of ancestral material in a state helps to break ties by assigning a smaller score to simpler configurations.

The method of computing the lower bound *L* depends on the complexity of the dataset, with a trade-off between accuracy and computational cost. For relatively small datasets, it is feasible to compute *R*_*min*_ exactly using Beagle. *HB* refers to the haplotype bound, employing the improvements afforded by first calculating local bounds for incompatible intervals, and applying a composition method to obtain a global bound (Myers and Griffiths, 2003). *HK* refers to the Hudson-Kaplan bound (Hudson and Kaplan, 1985); this is quick but less accurate, so is reserved for larger, more complex configurations. Note that these bounds are computed under the infinite sites assumption.

The particular form and components of the score were chosen through simulation testing; we found that the given formula provides a good level of informativeness regarding the quality of a possible state.

#### 2.2.3. Event cost

Each type of event is assigned a cost, which gives a relative measure of preference for each event type in the reconstructed history:

- *C*_*R*_: the cost of a single recombination event, defaults to 1.
- *C*_*RR*_: the cost of performing two successive recombinations, defaults to 2. It is sufficient to consider at most two consecutive recombination events before a coalescence (Lyngsø et al., 2005); this type of event also captures the effects of gene conversion.
- *C*_*RM*_: the cost of a recurrent mutation. If 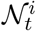 is formed from 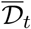 by a recurrent mutation in a column representing *k* agreeing sites, this corresponds to proposing *k* recurrent mutation events, so the cost is 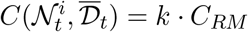.
- *C*_*SE*_: this event is a recurrent mutation which affects only one sequence in the original dataset, i.e. it occurs on the terminal branches of the ARG. Thus, the event can be either a regular recurrent mutation, or an artefact due to sequencing errors. The cost can be set to equal *C*_*RM*_, or lower if the presence of sequencing errors is considered likely.

KwARG allows the specification of a range of event costs as tuning parameters, as well as the number *Q* of independent runs of the algorithm to perform for each cost configuration. The proportions of recombinations to recurrent mutations in the solutions produced by KwARG can be controlled by varying the ratio of costs for the corresponding event types.

#### 2.2.4. Selection probability

The method of selecting the next state from a neighbourhood of candidates will impact on the efficiency and performance of the algorithm. At one extreme, selecting at random amongst the states will mean that the solution space is explored more fully, but will be prohibitively inefficient in terms of the number of runs needed to find a near-optimal solution. On the other hand, always greedily selecting the move with the minimal score will quickly identify a small set of solutions for each cost configuration, at the expense of placing our faith in the ability of the score to assess the quality of the candidate states accurately.

We propose a selection method that is intermediate between these two extremes, randomising the selection but focusing on moves with near-minimal scores. A pseudo-score for state 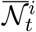 is calculated:

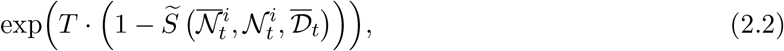

where

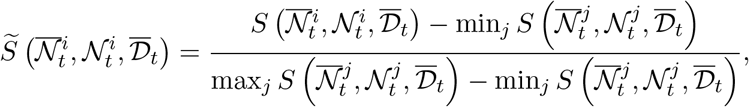

and states in 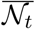 are selected with probability proportional to their pseudo-score. The annealing parameter *T* controls the extent of random exploration; *T* = 0 corresponds to choosing uniformly at random from the neighbourhood of candidates, and *T* = ∞ to always choosing a state with the minimal score. The default value of *T* = 30 was chosen following simulation testing, which showed that this provides a good balance between efficiency and thorough exploration of the neighbourhood.

### 2.3. Output

The default output consists of the number of recombinations and recurrent mutations in each identified solution; an example for the Kreitman dataset is given in Table 1. Each iteration is assigned a unique random seed, which can be used to reconstruct each particular solution and produce more detailed outputs, such as a detailed list of events in the history, the ARG in several graph formats, or the corresponding sequence of marginal trees.

**Table 1.** Example output of KwARG for the Kreitman dataset. SE: number of recurrent mutations occurring on terminal branches of the ARG (possible sequencing errors). RM: number of other recur-rent mutations. R: number of recombinations. Last column gives the total number of neighbourhood states considered.

| | Seed | $T$ | $C_{SE}$ | $C_{RM}$ | $C_R$ | $C_{RR}$ | SE | RM | R | $\sum_t \vert \mathcal{N}_t\vert $ |
| --- | --- | --- | --- | --- | --- | --- | --- | --- | --- | --- |
| 1 | 2263536315 | 30.0 | 100.00 | 100.00 | 1.00 | 2.00 | 0 | 0 | 7 | 143 |
| 2 | 2347021759 | 30.0 | 0.90 | 0.91 | 1.00 | 2.00 | 1 | 0 | 6 | 853 |
| 3 | 1791455164 | 30.0 | 0.80 | 0.81 | 1.00 | 2.00 | 1 | 0 | 5 | 728 |
| 4 | 1684879495 | 30.0 | 0.60 | 0.61 | 1.00 | 2.00 | 2 | 0 | 4 | 783 |
| 5 | 1884182000 | 30.0 | 0.40 | 0.41 | 1.00 | 2.00 | 3 | 0 | 3 | 806 |
| 6 | 1900122424 | 30.0 | 0.20 | 0.21 | 1.00 | 2.00 | 5 | 0 | 2 | 702 |
| 7 | 2111915557 | 30.0 | 0.10 | 0.11 | 1.00 | 2.00 | 8 | 0 | 1 | 833 |
| 8 | 2888657821 | 30.0 | 0.01 | 0.02 | 1.00 | 2.00 | 10 | 0 | 0 | 715 |

## 3. Performance on simulated data

### 3.1. Comparison to Beagle

Disallowing recurrent mutation, the accuracy of KwARG’s upper bound on *R*_*min*_ was tested by comparison with Beagle (Lyngsø et al., 2005). 1,100 datasets were simulated using msprime (Kelleher et al., 2016), under the infinite sites assumption (parameters: *N*_*e*_ = 1, mutation rate per generation per site 0.02, recombination rate per site 0.0003, 40 sequences of length 2,000bp). Of the generated datasets, 38 had no incompatible sites, and runs were terminated if Beagle took over 10 minutes to complete (which happened in 25 cases), leaving 1,037 datasets for testing. The parameters were chosen to produce datasets on which Beagle could be run within a reasonable amount of time; the value of *R*_*min*_ for the simulated datasets varied between 1 and 10.

Using the default annealing parameter *T* = 30, KwARG found *R*_*min*_ in all cases. In 97% of the runs, this took under 5 seconds of CPU time (on a 2.7GHz Intel Core i7 processor); all but one run took less than 40 seconds. In 93% of the runs, 1 iteration was sufficient to find an optimal solution; in 99% of the runs, 5 iterations were sufficient. Beagle found the exact solution in 5 seconds or less in 86% of cases; for datasets with a small *R*_*min*_ Beagle runs relatively quickly (median run time for *R*_*min*_ = 5 was 1 second, compared to KwARG’s 0.3 seconds). For more complex datasets, KwARG finds an optimal solution much faster; for *R*_*min*_ = 9, the median run time of Beagle was 56 seconds, compared to KwARG’s 3 seconds.

Setting *T* = 10 and *T* = ∞ resulted in 5 and 22 failures to find an optimal solution, respectively, when KwARG was run for *Q* = 1000 iterations per dataset (or terminated after 10 minutes have elapsed), demonstrating that setting the annealing parameters too low or too high results in deterioration of performance.

The left panel of Figure 3 illustrates the results, and shows the relationship between the true simulated number of recombinations and *R*_*min*_. This demonstrates that in many cases, substantially more recombinations have occurred than can be confidently detected from the data.

**Figure 3.**
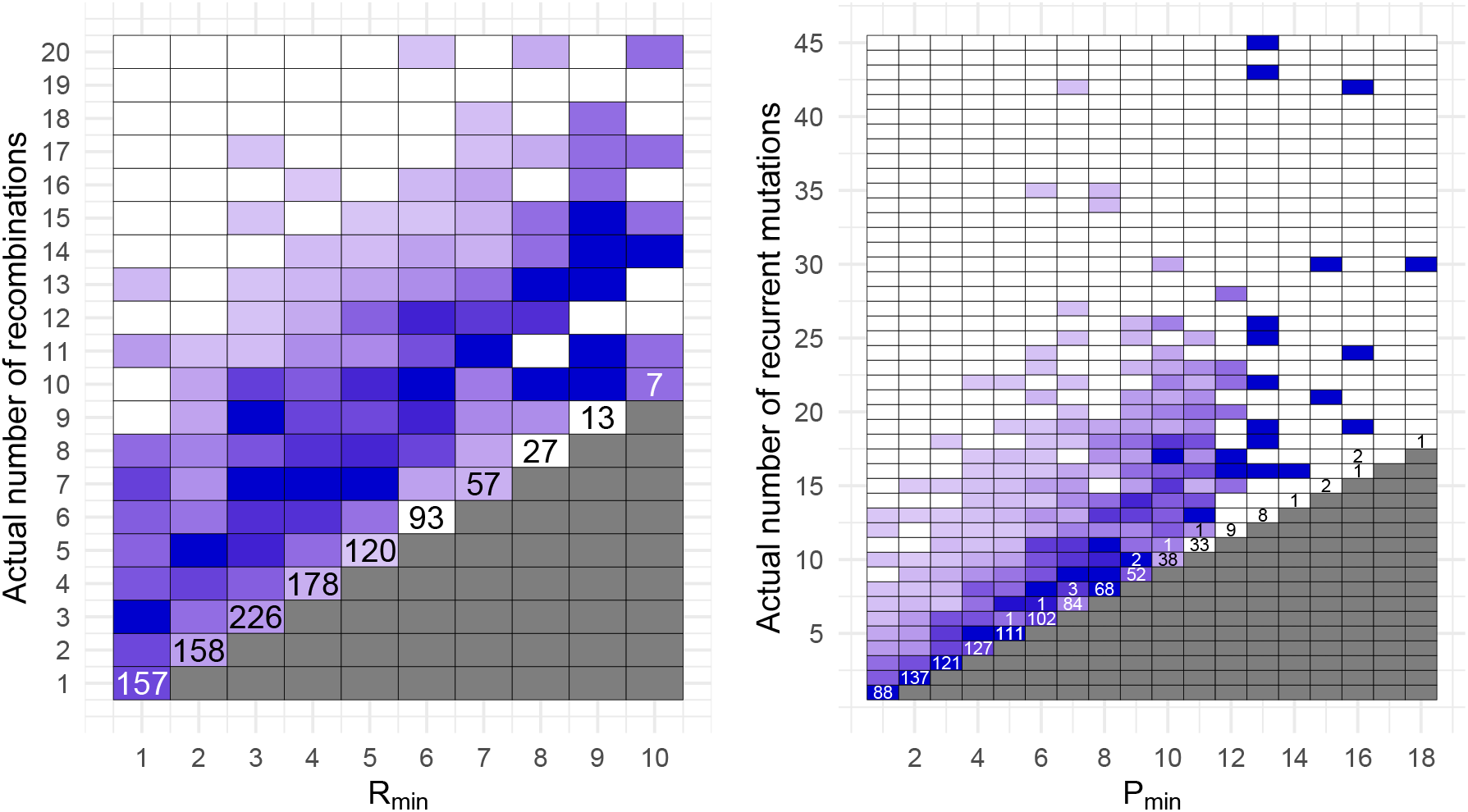
Left: number of simulated recombinations against *R*_*min*_. Right: number of simulated recurrent mutations against *P*_*min*_. Cell colouring intensity is proportional to the number of datasets generated for each pair of coordinates. Numbers in each cell correspond to the number of cases where for a dataset with the minimum number of events given on the *x*-axis, KwARG inferred the number of events given on the *y*-axis (unlabelled cells correspond to 0 such cases).

### 3.2. Comparison to PAUP*

Disallowing recombination, the quality of computed upper bounds on *P*_*min*_ was tested by comparison with PAUP* (Swofford, 2001, version 4.0a168), which was used to compute the exact minimum parsimony score via branch-and-bound. 1,100 genealogies were simulated using msprime (parameters: 20 sequences, *N*_*e*_ = 1). For each tree, Seq-Gen (Rambaut and Grass, 1997) was used to add mutations (parameters: 1,000 sites, mutation rate per generation per site set by the scaling constant *s* = 0.01); only transitions were allowed, to fulfil the requirement that sites mutate between exactly two states. 1,063 datasets exhibited incompatibilities caused by recurrent mutations. KwARG was run for a total of *Q* = 600 iterations per dataset; 150 of these were used to estimate *R*_*min*_, and 450 were run with a range of costs to estimate *P*_*min*_. The runs were terminated after 10 minutes (if 600 iterations had not been completed by then, the results were discarded; this happened in 69 cases).

KwARG failed to find *P*_*min*_ in 11 (1.1%) cases out of 994 successful runs. The results are illustrated in the right panel of Figure 3. Where KwARG failed to find an optimal solution, in all 11 cases it was off by just one recurrent mutation. Figure 3 also demonstrates that a substantial proportion of recurrent mutations do not create incompatibilities in the data, and the number of actual events often far exceeds *P*_*min*_.

### 3.3. Comparison to SHRUB and SHRUB-GC

The performance of KwARG on larger datasets was tested by comparison to SHRUB (Song et al., 2005) and SHRUB-GC (Song et al., 2006). Both methods implement a backwards-in-time construction of ARGs, using a dynamic programming approach to choose among possible recombination events. SHRUB produces an upper bound on *R*_*min*_ under the infinite sites assumption. SHRUB-GC also allows gene conversion events; setting the maximum gene conversion tract length to 1 makes this equivalent to recurrent mutation. The algorithm seeks to minimise the total number of events, essentially assigning equal costs to recombination and recurrent mutation. This differs from KwARG in that a single solution is produced for a given dataset, rather than a full range of solutions varying in the number of recombinations and recurrent mutations.

Using msprime and Seq-Gen, 300 datasets of 100 sequences were simulated, with a range of mutation and recombination rates and sequence lengths of 2,000, 5,000, 8,000 and 10,000 bp. For each dataset, KwARG was run for a total of *Q* = 260 iterations, with the default cost configurations and *T* = 30. The resulting upper bound on *R*_*min*_ was compared to that produced by SHRUB, and the minimum number of events over all identified solutions was compared to the solution produced by SHRUB-GC (configured to allow length-1 gene conversions).

KwARG obtained solutions at least as good as SHRUB’s in 292 (97.3%) of 300 cases, outperforming it in 35 (11.7%) instances. KwARG obtained solutions at least as good as SHRUB-GC in 296 (98.7%) cases, outperforming it in 2 instances. The results and the run times are illustrated in Figures 7 and 8 of Appendix C. On average, for relatively small and simple datasets, KwARG takes approximately the same time per one iteration as a run of SHRUB or SHRUB-GC, and outperforms both programs on more complex datasets.

## 4. Application to Kreitman data

The performance of KwARG is illustrated on the classic dataset of Kreitman (1983, Table 1); this is not close to the performance limit of KwARG, but has been widely used for benchmarking algorithms used for ARG reconstruction. The dataset consists of 11 sequences and 2,721 sites, of which 43 are polymorphic, of the alcohol dehydrogenase locus of *Drosophila melanogaster*. The data is shown in Figure 4, with columns containing singleton mutations removed for ease of viewing. Applying the ‘Clean’ algorithm, as described in Section 2.2.1, reduces this to matrix of 9 rows and 16 columns. KwARG was run with the default parameters, *Q* = 500 times for each of 13 default cost configurations given in Appendix B. An example of the output is shown in Table 1.

**Figure 4.**
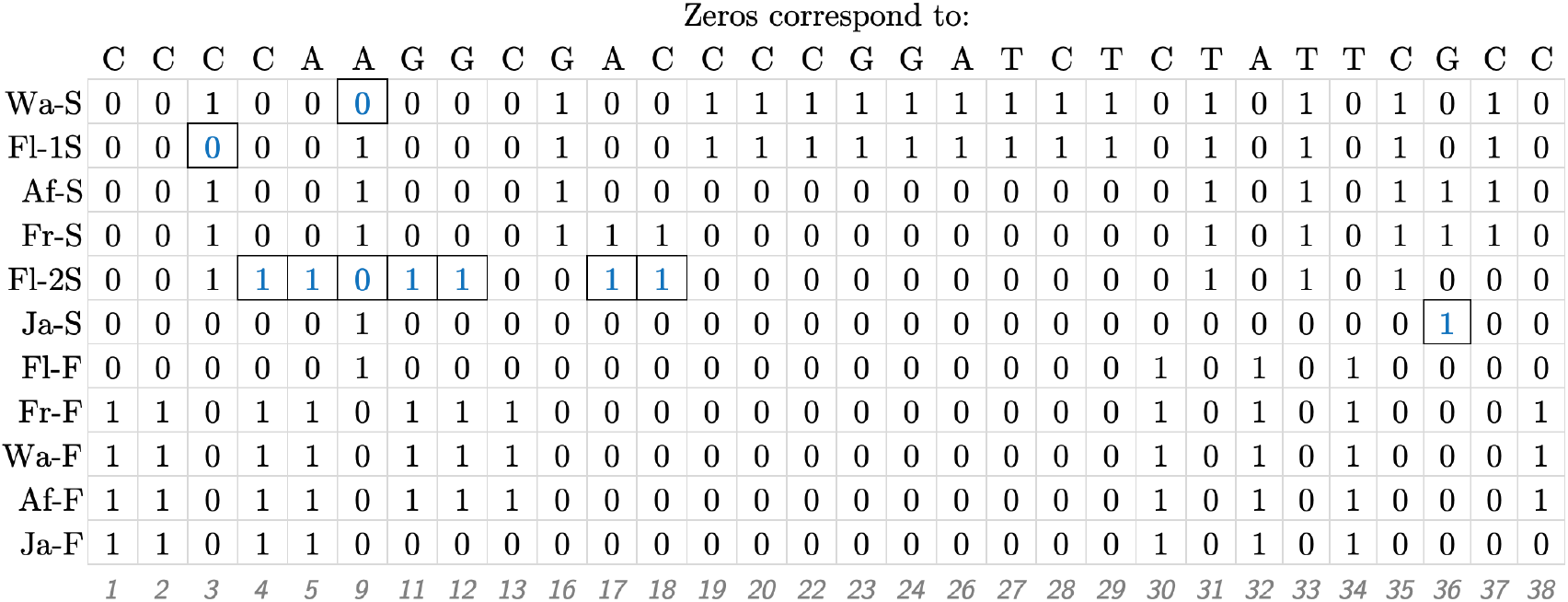
Illustration of the Kreitman dataset. The 11 sequences labelled as in Kreitman (1983); polymorphic sites are labelled 1–43 and columns with singleton mutations are not shown.

KwARG correctly identified the *R*_*min*_ of 7 and the *P*_*min*_ of 10 (confirmed by running Beagle and PAUP*, respectively). The 6,500 iterations of KwARG took just under 9 minutes to run. Of these, 1,829 (28%) resulted in optimal solutions; some are shown in Table 1. KwARG identified multiple combinations of recombinations and recurrent mutations that could have generated this dataset. By default, slightly cheaper costs are assigned to recurrent mutations if they happen on terminal branches, so the results show a bias towards solutions with more *SE* events for each given number of recombinations.

The ten recurrent mutations appearing in the solution in row 8 of Table 1 are highlighted on the dataset in Figure 4. It is striking that 7 of these 10 recurrent mutations affect the same sequence Fl-2S. In fact, these 7 recurrent mutations could be replaced by 3 recombination events affecting sequence Fl-2S, with breakpoints just after sites 3, 16, and 35; leaving the other identified recurrent mutations unchanged yields the solution in row 5 of Table 1. These findings suggest that the sequence may have been affected by cross-contamination or other errors during the sequencing process, or it could indeed be a recombinant mosaic of four other sequences in the sample. This recovers the results obtained by Stephens and Nei (1985), who posited the recombinant origins of sequence Fl-2S following manual examination of a reconstructed maximum parsimony tree, which also highlighted the five consecutive mutations identified by KwARG. The ARG corresponding to the solution in row 5 of Table 1, visualised using Graphviz (Ellson et al., 2004), is shown in Figure 5.

**Figure 5.**
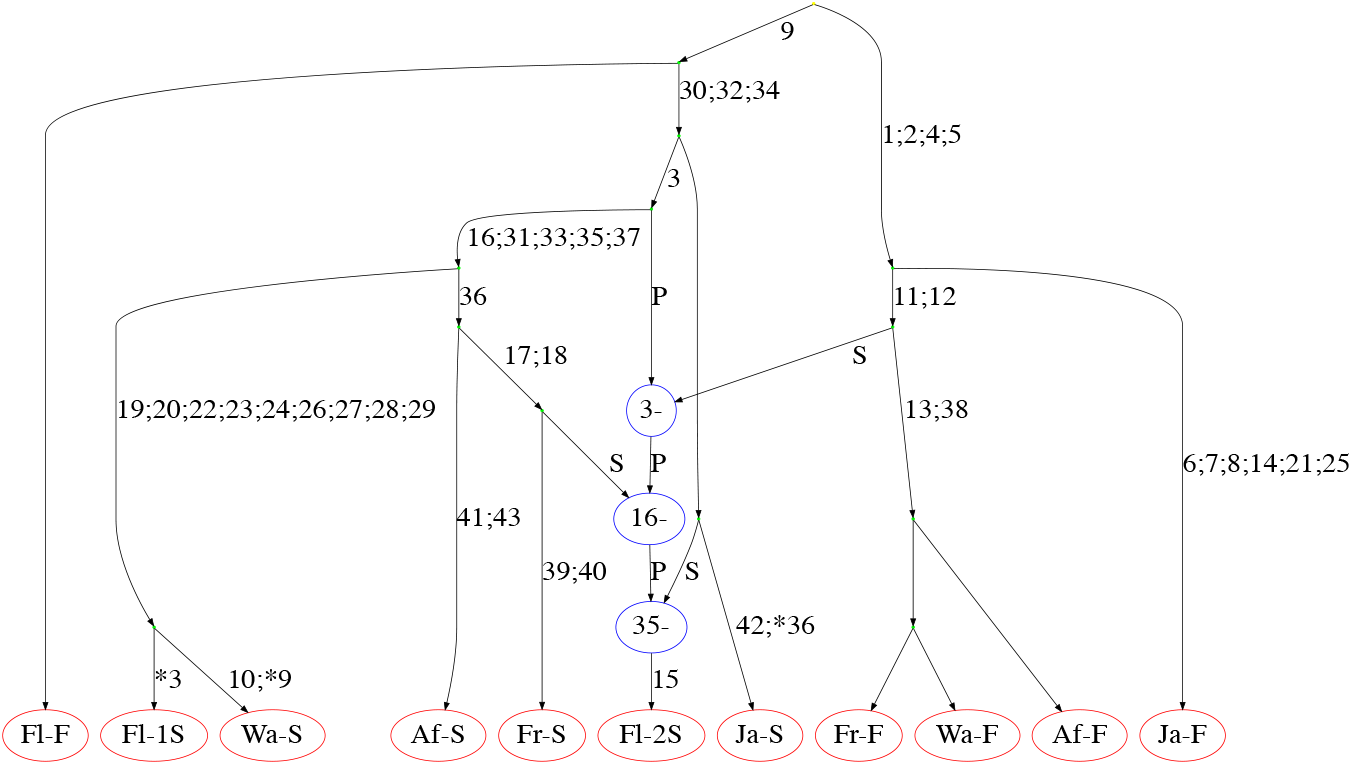
ARG constructed for the Kreitman data. Edges are labelled with sites undergoing mutations; recurrent mutations are prefixed with an asterisk. Recombination nodes, in blue, are labelled with the recombination breakpoint; material to the right (left) of the breakpoint is inherited from the parent connected by the edge labelled *S* (*P*) for “suffix” (“prefix”).

Examination of the identified solutions also shows that site 36 of sequence Ja-S “necessitates” two of the seven recombinations inferred in the minimal solution in the absence of recurrent mutation, while sites 3 and 9 in sequences Wa-S and Fl-1S, respectively, each create incompatibilities that could be resolved by one recombination.

## 5. Discussion

Methods for the reconstruction of parsimonious ARGs generally rely on the infinite sites assumption. When examining the output ARGs, it is often difficult to tell how much the inferred recombination events actually affect the recombining sequences. As is the case with the Kreitman dataset, sometimes further examination reveals that two crossover recombination events have the same effect as one recurrent mutation, raising questions about which version of events is more likely. KwARG removes the need for such manual examination, and provides an automated way of highlighting such cases, which is particularly useful for larger datasets.

The solutions identified by KwARG differ in the proportion of recurrent mutations to recombinations, ranging from an explanation that invokes only recombination events to one that invokes only mutation events. Quantifying the likelihood of each scenario will be application-specific; for instance, one can choose a reasonable model of evolution for the population being studied, and identify the most likely solution under a range of reasonable mutation and recombination rates. When the presence or absence of recombination is not certain, then should the number of recurrent mutations needed to explain the dataset be infeasibly large, this provides evidence for the presence of recombination; this is the idea underlying the homoplasy test of Maynard Smith and Smith (1998). If the largest “reasonable” number of recurrent mutations is then estimated, KwARG can be used to say how many additional recombination events are required to explain the dataset.

KwARG performs well when compared against exact methods for the ‘recombination-only’ and ‘mutation-only’ scenarios. Because of the random exploration incorporated within KwARG, it should be run multiple times on the same dataset before selecting the best solutions; the optimal run length of KwARG will be constrained by timing and the available computational resources. To gauge whether KwARG has run enough iterations, one could proceed by calculating *R*_*min*_ and *P*_*min*_ either exactly (if the data is reasonably small) or using other heuristics-based methods (such as SHRUB or PAUP*), to confirm whether KwARG has found good solutions at these two extremes.

Further improvements could be obtained by amending the calculation of lower bounds within the cost function in order to account for the presence of recurrent mutation. Other avenues for further work include explicitly incorporating gene conversion as a possible type of recombination event with a separate cost parameter, with a view to developing the underlying model of evolution to even more closely reflect biological reality.

## 6. Acknowledgements

This work was supported by the OxWaSP CDT under the EPSRC and MRC grant EP/L016710/1, and by the Alan Turing Institute under the EPSRC grant EP/N510129/1.

## Appendix A. KwARG pseudocode

Let 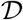 be an input data matrix with entries 0, 1 or ★. Denote by 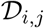 the entry of 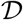 at position (*i, j*). Let 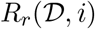 and 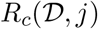 denote the resulting matrix when the *i*-th row or the *j*-th column of 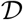 is deleted, respectively. Let the history 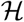 be a set storing all of the intermediate states visited on the path from 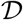 to the root of the ARG.

**Algorithm 1:**
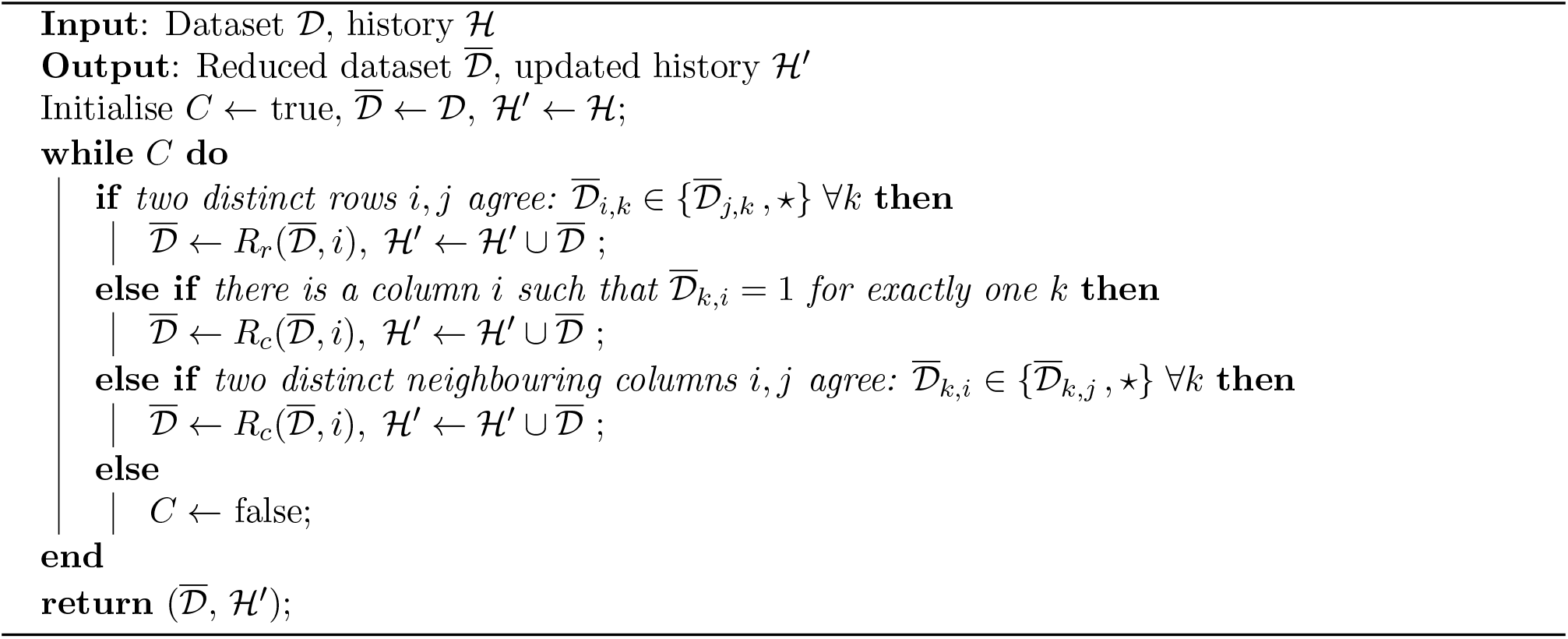
Clean (adapted from Song and Hein, 2003)

Define the following operations:

1. Recurrent mutation: 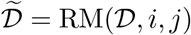 is the result of a recurrent mutation in row *i* at column *j*; 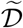 is obtained from 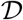 by changing the (*i, j*)-th entry from 0 to 1 or from 1 to 0.
2. Recombination: 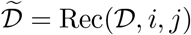 is the result of a recombination in row *i* with breakpoint just after column *j*. Namely, 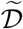 is obtained from 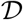 by inserting a copy of the *i*-th row just below itself, and setting 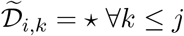 and 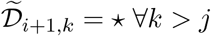.
3. Two consecutive recombinations: 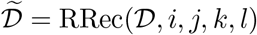 is the result of performing two recombinations, in rows *i* and *k* with breakpoints at *j* and *l*, respectively.

Note that for recombination events, not all row and column positions should to be considered, as some moves are guaranteed not to resolve any incompatibilities in the dataset. We apply the ideas detailed in Lyngsø et al. (2005, Section 3.3) to restrict the rows and breakpoints considered for recombination events. Suppose that as a result, 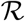 is the list of row and column indices (*i, j*) to consider for recombination events, and 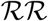 is the list of indices (*i, j, k, l*) to consider for two consecutive recombination events.

**Algorithm 2:**
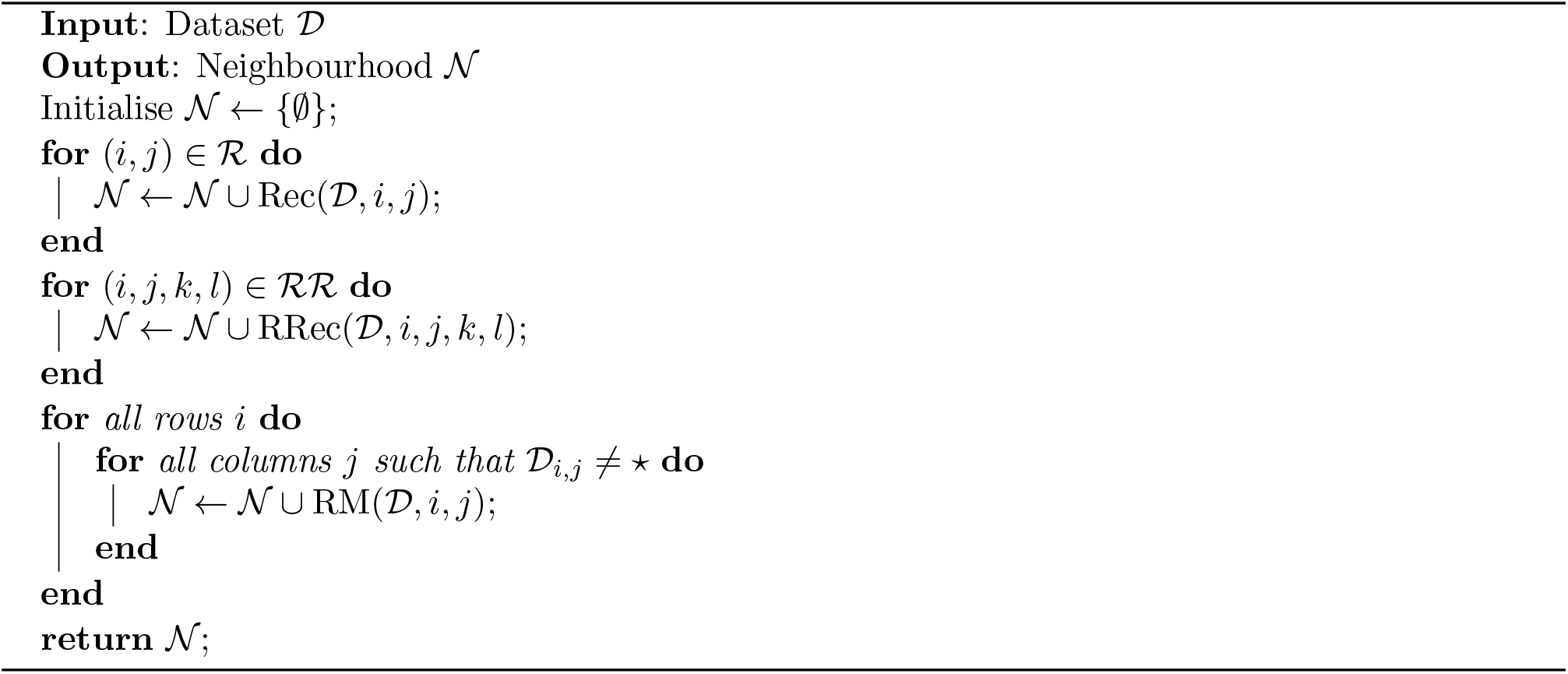
Neighbourhood

**Algorithm 3:**
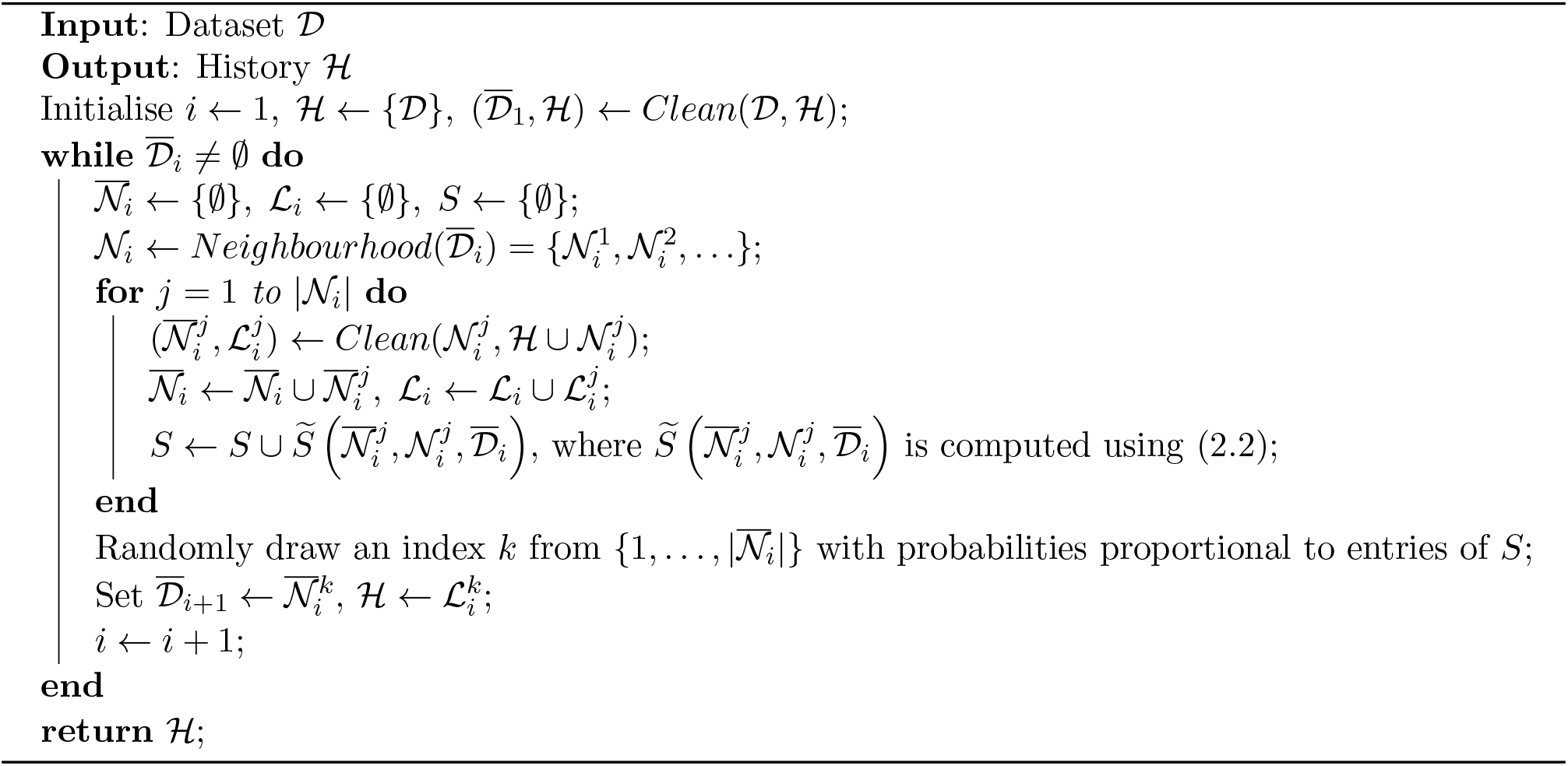
KwARG

## Appendix B. Default cost configuration

If the number of iterations *Q* > 1 is specified but no costs are input, KwARG runs each of the following 13 cost configurations *Q* times:

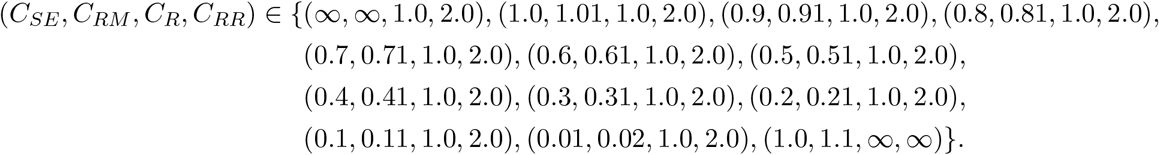

The effectiveness of this is illustrated in Figure 6, which is based on the set of all possible minimal solutions identified for the Kreitman dataset. Fixing *C*_*R*_ = 1.0 and *C*_*RR*_ = 2.0, each tile represents a pair (*C*_*SE*_, *C*_*RM*_). Each tile is coloured and labelled according to the corresponding cost-optimal solution, in the form {*x, y, z*}, giving the number of *SE*, *RM* and recombination events, respectively. For instance, if *C*_*SE*_ = 0.5 and *C*_*RM*_ = 0.61, the solutions {3, 0, 3} (with cost 3 · 0.5 + 3 · 1.0 = 4.5) and {5, 0, 2} (with cost 5 · 0.5 + 2 · 1.0 = 4.5) have the lowest costs over all feasible solutions.

**Figure 6.**
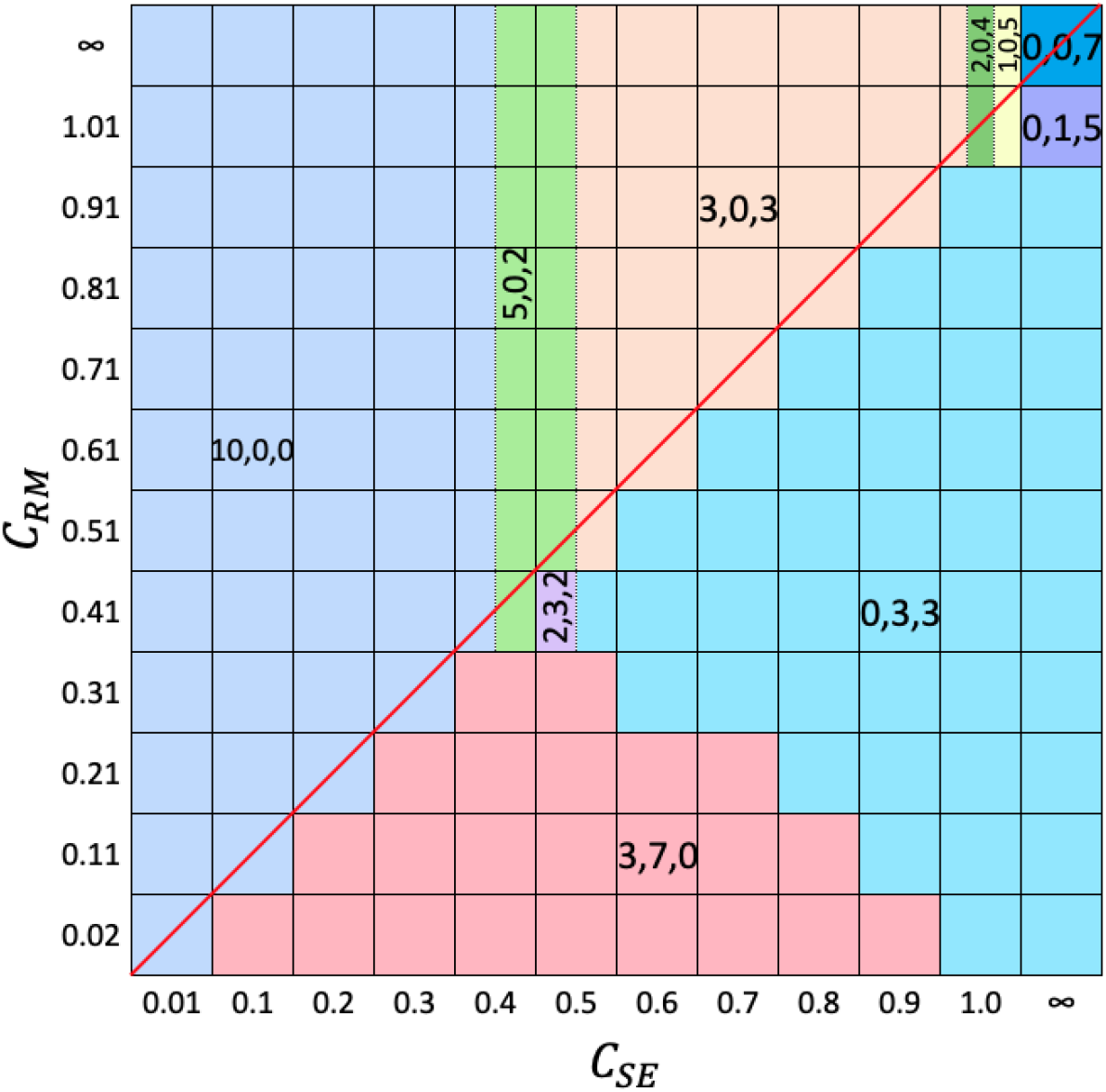
Solution tile plot for the Kreitman dataset.

The default cost configuration includes all pairs (*C*_*SE*_, *C*_*RM*_) on the diagonal in this plot, falling on the red line. This line crosses all optimal solutions which maximise the number of *SE* events for each possible number of recombinations. Such events affect only a single sequence at a single site in the input dataset, so are, in a sense, more parsimonious than recurrent mutations occurring on internal branches.

## Appendix C. Results of comparison to SHRUB and SHRUB-GC

**Figure 7.**
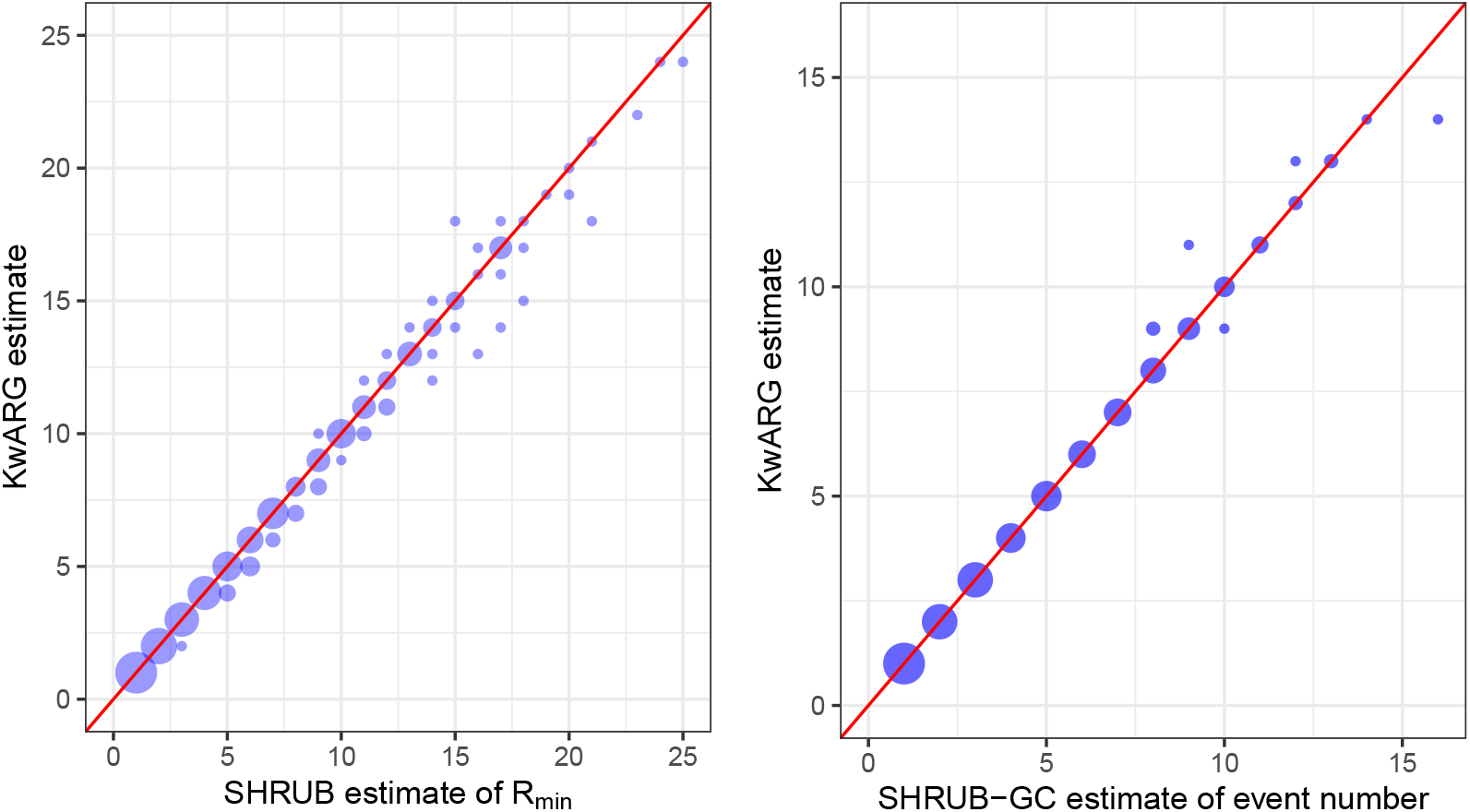
Comparison of KwARG to SHRUB and SHRUB-GC. *x*-axis: estimate produced by SHRUB (left) and SHRUB-GC (right). *y*-axis: estimate produced by KwARG. Instances where equally good solutions were found lie on the red diagonal line. Size of points is proportional to the number of corresponding datasets.

**Figure 8.**
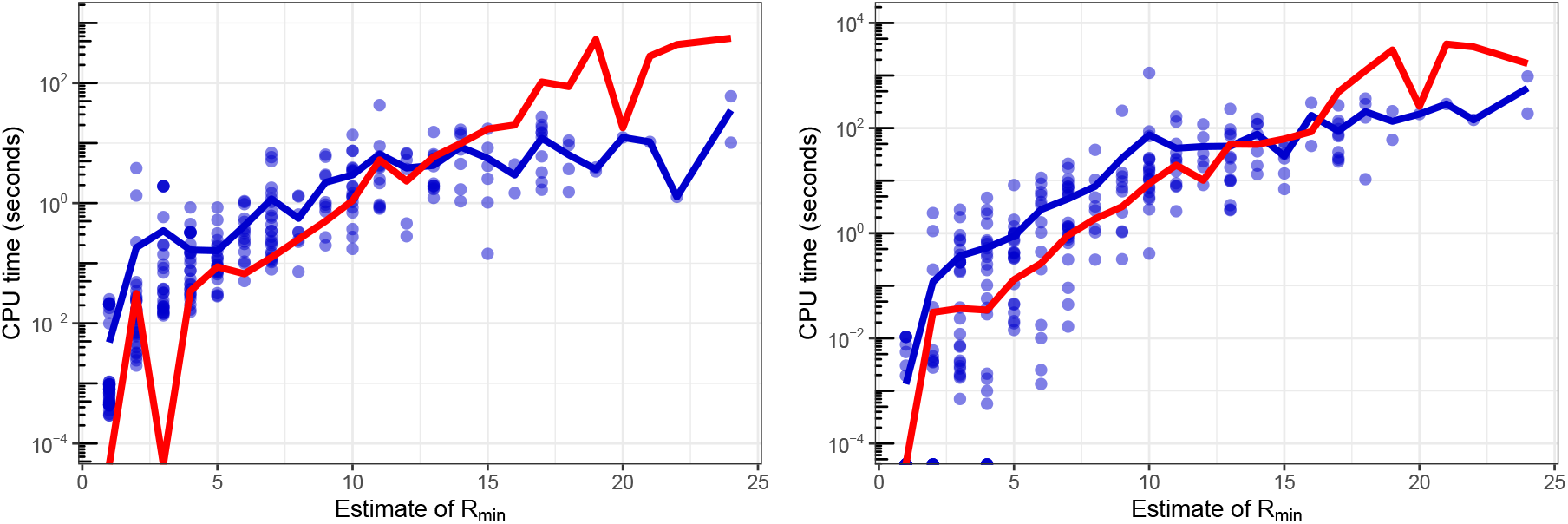
Blue points: time taken to run *Q* = 20 iterations of KwARG (left: disallowing recurrent mutations, right: allowing both recombination and recurrent mutation). Blue lines: mean values. Red line: mean run time of SHRUB (left) and SHRUB-GC (right). Time in seconds is given on a log scale.

